# Retrospective analysis of *Pseudomonas aeruginosa* clinical isolates for secreted PrpL protease: a key virulence factor associated with corneal tissue damage

**DOI:** 10.64898/2025.12.11.693724

**Authors:** Douglas S. Parker, Onyedikachi C. Azuama, Kashaf Zafar, Jon D. Goguen, John M. Leong, Nikhat Parveen

## Abstract

**Background:** *Pseudomonas aeruginosa* is a ubiquitous organism that adapts well in different environments. It is an opportunistic bacterial pathogen that produces a wide range of virulence factors, colonizes lungs to cause pneumonia, causes non-healing wounds especially in burn victims, and is a major culprit in destructive keratitis. It can reach the cornea through reusable, extended-use contact lenses and by contaminated eyedrops and artificial tears. Secreted proteases of *P. aeruginosa* together with pyocyanin metabolite, which inhibits Serine Protease Inhibitors (Serpins) activity contribute to severe tissue damage during infection.

**Methods:** *P. aeruginosa* strains isolated from different clinical sites were obtained from different researchers and clinicians for this study. We examined *P. aeruginosa* strains and secretion defective and prpL knockout mutants in PA64481 strain for lysyl endopeptidase (PrpL, a serine protease that cleaves after a lysine residue) activity using serine protease specific D-Val-Leu-Lys-p-nitroanalide substrate. We also determined pyocyanin production in these strains.

**Results:** Examination of secreted milieu from *P. aeruginosa* showed that 41 corneal isolates had detectable lysyl endopeptidase activity associated with PrpL at levels significantly higher than by 27 non-corneal isolates. We found that PrpL is secreted by the *xcp*-based type II secretion system. Bacterial culture supernatants displaying higher PrpL activity and not low secreted PrpL levels disrupted corneal epithelial cell monolayers *in vitro*, which is consistent with a role of this protease in destructive keratitis. Many examined strains also produced high levels of pyocyanin.

**Conclusions:** This retrospective examination of clinical *P. aeruginosa* suggests that high levels of PrpL and pyocyanin-producing isolates are more prevalent among corneal isolates and could enhance tissue damage during infection. Supporting this premise, corneal epithelial cell monolayers disrupted by high PrpL-producing strains but remained intact after treatment with *P. aeruginosa* mutants culture supernatants that lack or have reduced secreted PrpL.

## INTRODUCTION

*Pseudomonas aeruginosa* is a ubiquitously existing Gram-negative bacterium which is also an important human pathogen. At least one strain of *P. aeruginosa* can produce lethal infections in mice, flies, nematodes, and plants, making it the pathogen *par excellence* [1]. In humans, it is an opportunist that requires tissue damage and/or weakened host defenses to produce significant disease. Consistent with its adaptation to parasitism, *P. aeruginosa* utilizes type III secretion system to directly inject protein effectors into the cytoplasm of eukaryotic cells and employs other secretion systems to export several virulence factors into the environment. *P. aeruginosa* infection often causes non-healing wounds especially in burn victims and inflicts acute to chronic lung disease in the intensive care units and among cystic fibrosis patients. It is a primary contributor in corneal infections and ulcerative keratitis particularly among extended-wear contact lens users [2–4]. Furthermore, immunocompromised and post-cataract surgery individuals are predisposed to *P. aeruginosa* infection [5]. Emergence of multidrug-resistant (MDR) and extensively drug-resistant (XDR) *P. aeruginosa* strains has greatly complicated treatment of these infections. A recent outbreak of MDR/XDR *P. aeruginosa* corneal infections due to contaminated artificial tear eye drops indicates that this infection can result in blindness and even death due to sepsis [6–8], thus needing better understanding of the critical virulence factors of *P. aeruginosa*.

*P. aeruginosa* secretes several degradative proteases into the extracellular environment [9, 10]. Among these, alkaline protease, elastase A, and elastase B are firmly implicated in *P. aeruginosa* pathogenesis in cystic fibrosis patients [11] but not in keratitis [12]. Mammalian protease inhibitors regulate the activity of host, and several *P. aeruginosa* proteases, potentially minimizing tissue injury they cause. The fourth, a ∼27 kD serine protease labeled as Protease IV [13, 14] was previously purified from PA103-29 strain and implicated in ulcerative keratitis in an experimental animal model [13–19]. This serine protease/PrpL is highly specific for peptide bonds at the carbonyl group of lysine residue (lysyl-endopeptidase) in protein, ester, or amide substrates. Therefore, D-Val-Leu-Lys-p-nitroanalide or S2251 chromogenic substrate is ideal for PrpL activity determination.

Complex mechanisms regulate the expression of different virulence factors of *P. aeruginosa*. PvdS (PA2426 in PA01 strain) is an iron-starvation regulator that drives the expression of genes involved in the synthesis of the pyoverdine siderophore [20] and extracellular protease PrpL (PvdS-regulated endopeptidase, lysyl class)/Protease IV and the secreted exotoxin A [21]. Activities of serine proteases are regulated in hosts largely through production of a group of protease inhibitors, such as the serpins (<u>Ser</u>ine <u>p</u>rotease <u>in</u>hibitors) class [22, 23]. Among the most prominent anti-proteases is α1-protease inhibitor (α1-PI), which forms an enzymatically inactive complex with various serine proteases, may control host proteases [23, 24], they could similarly limit the damage caused by bacterial serine proteases, such as PrpL. Interestingly, proteases also play critical roles in *P. aeruginosa* competitiveness and fitness in hosts by cleaving host serpins, enhancing shedding of host proteins to increase invasiveness and colonization in cornea and airway in cystic fibrosis patients [25, 26].

*P. aeruginosa* has adapted to the potentially antagonistic effects of host protease inhibitors like α1-PI on bacterial proteases by also secreting pyocyanin, a phenazine compound that counteracts host protease inhibitors. Oxidation of methionine at the active site of α1-PI results in inactivation of this inhibitor [27]. Pyocyanin stimulates production of oxidants such as superoxide (O2^.-^), and hydrogen peroxide (H_2_O_2_) under aerobic conditions and in the presence of reducing compounds such as NADH, it could also control the intracellular redox homeostasis in bacteria [28] and hosts. Pyocyanin-treated α1-PI failed to inactivate the serine proteases and the inhibitor was then degraded by either trypsin or porcine pancreatic elastase *in vitro* [29]. Thus, pyocyanin and PrpL could synergistically exacerbate tissue damage by *P. aeruginosa* especially during corneal infection.

In this study, we show that consistent with a role in keratitis, PrpL activity in human corneal *P. aeruginosa* isolates was significantly (p=0.035) higher than that in non-corneal clinical isolates and PrpL was associated with the destruction of cultured corneal epithelial cell monolayers not observed in *xcpA* and *prpL* mutants. Based upon literature and our results here, we anticipate that pyocyanin, an antagonist of serpins, would augment the activity of PrpL and exacerbate tissue destruction, highlighting the importance of proteolytic damage in the pathogenesis of *P. aeruginosa* in humans.

### Ethics Statement

Patients personal information, clinical details and other information associated with *P. aeruginosa* strains were not provided except documentation of the original isolation site. Therefore, informed consent from patients is not applicable for this study. However, corresponding author has approved exempt Institutional Protocol Board protocol to conduct studies with clinical bacterial isolates.

## Materials and Methods

### Bacterial strains, corneal cell line and growth conditions

Several *P. aeruginosa* strains including PA01 and PA103 were generously provided by Dara Frank. Other strains, including PA64481, were gifts of Darlene Miller of Bascom Palmer Eye Institute and Brenda Torres of the University of Massachusetts-Memorial clinical laboratories.

*P. aeruginosa* cultures were grown in iron sufficient Luria Bertani (LB) or in SM9 minimal medium containing 2 g Glucose, 3 g KH_2_PO_4_, 6 g Na_2_HPO_4_, 1g NH_4_Cl, 5 g NaCl, 0.2 μM CaCl_2_, 1 μM MgSO_4_ and 1mg Thiamine per liter, pH 6.9. Human corneal epithelial cell line developed by Araki-Sasaki by transformation with SV40 vector (HCE-T), was obtained from RIKEN, Japan. The HCE-T cell line was cultured in the medium containing DMEM: Ham12 in a 1:1 ratio, 5%FBS, 5µg/ml insulin, 0.1µg/ml cholera toxin, 10ng/ml hEGF, and 0.5%DMSO at 37°C in 5% CO2 incubator.

### Analysis of PrpL protease activity and determine levels of pyocyanin produced

We assayed PrpL protease (PA4175) activity using chromogenic plasmin substrate D-Val-Leu-Lys -p-nitroanalide or S2251 (Sigma Chemical Co.) and carried out the kinetic studies in microtiter plates at 37°C in a 100μl volume with 85 μl buffer, containing 50 mM Tris, pH 7.56, 0.1% Tween 80, 0.02% Sodium azide, 1% glycerol, 5 μl culture supernatant and 10 μl substrate (5 mM final) per well. The reproducibility of the experiment was ensured by performing each assay in triplicate and repeating the experiment at least three times. Total pyocyanin levels in the culture supernatants were extracted with acid, followed by chloroform, and determined colorimetrically as described previously [1].

### Purification of PrpL protease and raising antibodies

Ion exchange chromatography (DEAE-Sepharose) followed by size exclusion chromatography of a crude supernatant from a high PrpL-producing strain, PA64481 grown in minimal medium, yielded a single peak of activity with an estimated molecular mass of ∼27 kD. No other fraction showed significant PrpL activity. We used the pulverized PrpL protein band from PA64481 supernatant resolved in SDS-PAGE to raise polyclonal antibodies in mice.

### Mutagenesis and cloning of mutated DNA fragment

Random transposon mutagenesis of PA64481 was achieved by conjugational transfer of non-replicative plasmid pRK2013 containing Tn5 with gentamycin (Tn5Gm) from *E. coli* strain κ1849(pRK2013: Tn5Gm), which is unable to synthesize diaminopimelic acid (DAP^-^). We selected mutant on casein overlay on LBGent^75^ plates lacking DAP and selected transposon-insertion mutant colony that lacked a zone of casein digestion. The site of transposon insertion in the mutant was determined by cloning the fragment in pGATA (Amp^r^) vector followed by sequence analysis. We conducted directed mutagenesis of *prpL* gene by amplifying the gene and flanking region by PCR and disrupted gene with Gentamycin cassette was used to obtain the mutant in PA64481 strain by selecting knockout mutant (δPrpL) on antibiotic containing plate.

### Effect of *P. aeruginosa* culture supernatants on HCE-T cell line

The HCE-T monolayers on coverslips were incubated with culture supernatant diluted 1:1 in the cell culture medium up to 1h at 15 minutes interval, fixed with 3% paraformaldehyde prepared in phosphate buffered saline, and permeabilized with 0.1% Triton-X100 for 5 minutes. After washing three times with PBS at 5-minute intervals, the cells were stained to detect cytokeratins by incubating with 1:100 dilution of anti-keratin guinea pig antibodies (Sigma Chemical, Inc.) for 1h. After washing, coverslips were incubated with anti-guinea pig FITC (Sigma F6261) secondary antibodies together with TRITC-phalloidin (0.5µg/ml) for 1h for visualization of cytokeratins and filamentous actin, respectively. After mounting coverslips, we observed the slides with Nikon inverted fluorescence microscope.

## RESULTS

### Supernatants from *P. aeruginosa* corneal isolates exhibit higher PrpL activity

We quantitated PrpL lysyl endopeptidase activity in 41 corneal and 27 non-corneal *Pseudomonas* clinical isolates using the chromogenic substrate S2251. Although activity as high as 320 IU/ml was detected in the supernatants of a few non-corneal isolates, half of the isolates expressed negligible activity (Figure 1). We used PA01 strain, a wound isolate and first sequenced *P. aeruginosa* strain, as the Type strain in this study. Interestingly, higher pyocyanin levels than that of PA01 (OD_520nm_ = 0.9) were also detected in the majority of corneal and not most non-corneal isolated (Table 1). Furthermore, PrpL activity with the median OD_405_ of 2,800 among corneal strains (Figure 1) was significantly greater than that of non-corneal isolates (1,600, p=0.035). Thus, while PrpL activity may not be essential for establishment of corneal infection, association of PrpL production levels with corneal infecting isolates suggests that the protease contributes to destructive keratitis by *P. aeruginosa* in infected individuals.

**Figure 1.**
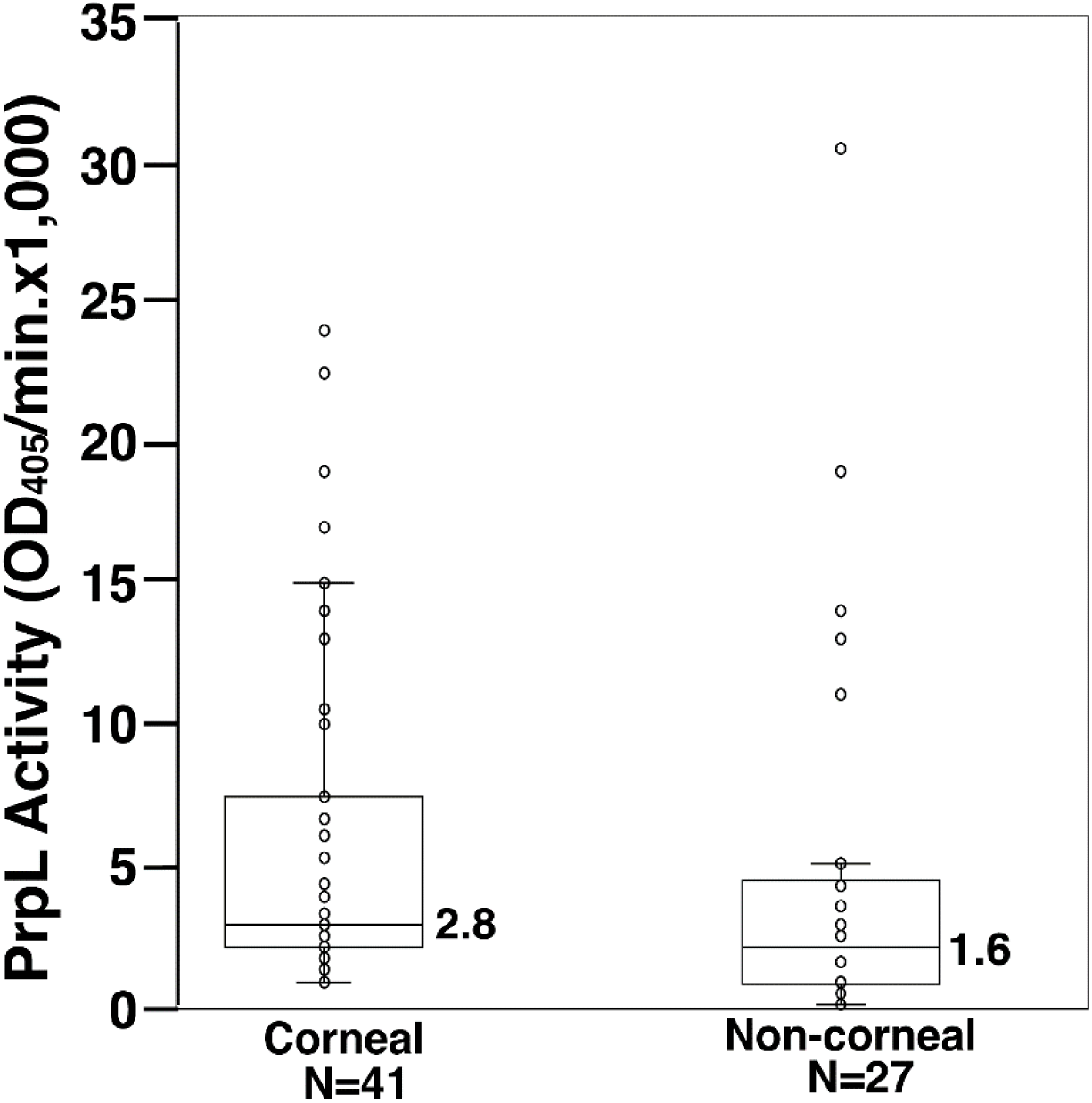
PrpL producing *P. aeruginosa* strains are over-represented in corneal infections. Kinetic analysis of lysyl endopeptidase activity in the culture supernatants from sixty-eight *P. aeruginosa* clinical isolates was determined using S2251 substrate with median OD_405_ value of in corneal (2.8×1,000) versus in non-corneal clinical (1.6×1,000) isolates. Mann-Whitney two-tailed U test for asymmetrically distributed data indicated a statistically significant (p=0.035) higher PrpL activity in corneal isolates.

**Table 1.** Grouping of *P. aeruginosa* strains based upon PrpL activity and pyocyanin levels.

| *P. aeruginosa* strains | Number of strains | Total pyocyanin (OD520≤0.9*) | Total pyocyanin (OD520>0.9) |
| --- | --- | --- | --- |
| PrpL Vmax per ml (>0.5 to<100) |  |  |  |
| (a) Corneal | 15 | 4 | 11 |
| (b) Non-corneal | 16 | 11 | 5 |
| PrpL Vmax per ml (100 to 500) |  |  |  |
| (a) Corneal | 10 | 1 | 9 |
| (b) Non-corneal | 7 | 3 | 4 |
| PrpL Vmax per ml (>500) |  |  |  |
| (a) Corneal | 16 | 2 | 14 |
| (b) Non-corneal | 4 | 0 | 4 |
| **Total** | 68 | 21 | 47 |

\* Pyocyanin level in PA01

### PA64481 strain secretes higher levels of active PrpL protease

We included the PA01 as well as PA103 in all experiments because PA103 demonstrate almost undetectable PrpL activity. PA103 strain was isolated from patient’s sputum and was originally considered to lack the secreted proteolytic activity; however, its one derivative PA103-29 exhibited intermediate PrpL activity [30]. Silver-stained SDS-PAGE gel of culture supernatants revealed a species with an apparent molecular mass of ∼27 kD, i.e., the predicted molecular mass of PrpL, that was much more prominent in PA64481 than in PA01 and PA103 (Figure 2A). This finding was reinforced by immunoblotting the three culture supernatants for PrpL (Figure 2B). Interestingly, pronounced PrpL activity in PA64481 resulted in autocleavage of the protease to some extent (Figures 2A and 2B). Overall, fewer proteins were observed in the culture supernatant of PA64481 (Figure 2A) likely due to the cleavage of many secreted proteins by highly active PrpL protease.

**Figure 2.**
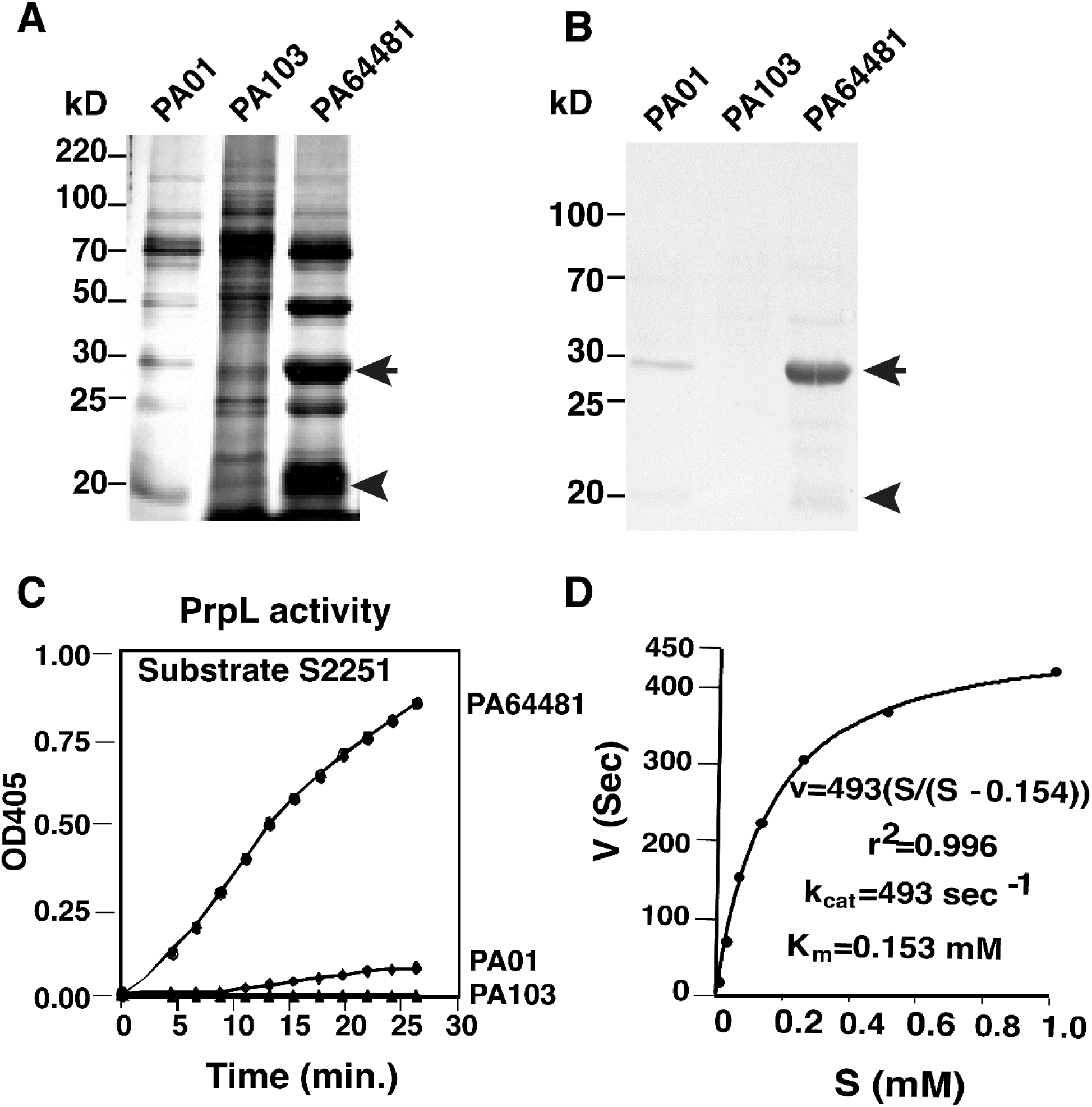
Detection of high specific activity of PrpL in PA64481 strain supernatant. **(A)** High secreted PrpL levels in PA64481, moderate level in PA01 were detected while PrpL was undetectable in PA103 strain. **(B)** Immunoblotting using anti-PrpL antiserum detected a major ∼27 kD PrpL band (arrow) and a minor band due to autocleavage of the PrpL protease (arrowhead). **(C)** Kinetic analysis of lysyl endopeptidase activity using S2251 substrate confirmed that secreted PrpL activity in PA64481 is significantly higher than PA01. No activity was detected in 30 minutes in PA103 culture supernatant. **(D)** K_cat_ and K_m_ of PrpL in PA64481 supernatant purified by Ion-exchange and size exclusion chromatography were determined. K_m_ of secreted PrpL was calculated to be 0.153 mM.

Bacterial supernatants of the wild-type PA103 displayed no PrpL protease activity after 30 minutes of incubation with the substrate while PA01 showed relatively low activity (Figure 2C), suggesting that these strains secrete only low levels of this protease. Therefore, PA64481 culture supernatant was subjected to chromatographic techniques described previously for the purification of PrpL from strain PA103 [13], and majority of activity was detected only in the ∼27 kD species. Sequence determination of the five N-terminal residues (AGYRD) of this purified protein was identical to the N-terminus of previously reported mature PrpL protease ([21], supplementary information). Using BCA kit, we estimated the concentration of purified PrpL in the SM9 PA64481 supernatant to be 150 ng of protease/ml, while estimated 0.4 mg of total protein was recovered previously after purification from 46 liters of PA103-29 culture supernatant [13]. The k_cat_ and K_M_ of PrpL secreted from PA64481, determined using the substrate S2251, were 493 µM sec^-1^ and 153 µM, respectively (Figs. 2D). In contrast, the V_max_ and K_m_ of this protease from PA103-29 (an *exoA* derivative of PA103 that secretes slightly higher levels of PrpL than the parental PA103 strain) were 0.74 µM/min and 727 µM, respectively [13]. Thus, more PrpL is produced and secreted by PA64481 than by PA103-29 or PA01. Higher k_cat_ of PA64481 PrpL was observed.

Comparison of deduced amino acid PrpL sequences demonstrated that whole protein sequences as well as the active site residues, serine, aspartate and histidine are conserved in all these serine proteases (Supplementary Figure 1A). Thus, activity in the culture supernatant is mainly reflective of PrpL secreted by each strain. Comparison with lysyl- and arginal endopeptidases showed that PrpL surprisingly showed slightly higher homology to the arginal endopeptidase LeR than to the lysyl endopeptidases AlK and LeK (Supplementary Figure 1B).

### Disruption of cultured corneal epithelial cells is associated with the level of secreted PrpL

To understand overall damage caused by PrpL during infection while serpins are inactivated by pyocyanin, we also determined the levels of this pigment in all strains. We detected higher pyocyanin levels in PA64481supernatant (OD_520_=1.7) while levels of this molecule were low to undetectable in PA01 and PA103 strains (OD_520_ 0.9 and 0.2, respectively). We added culture supernatants from three strains to the cultured HCE-T corneal epithelial monolayers. To eliminate the variation due to iron limiting conditions, LB grown bacterial culture supernatants were used to examine the effect of constitutively expressed and secreted PrpL protease. PA64481 supernatant destroyed HCE-T monolayers in less than 15 minutes (Figure 3, right panel). The cytokeratin organization and filamentous actin network were both disrupted by this treatment. Integrity of HCE-T cell monolayers was unaffected even after 1 h treatment with \PA01 and PA103 filter sterilized supernatants (Figure 3, left and central panels) supporting role of PrpL in corneal damage reported previously. Poor cytokeratin/phalloidin staining suggests that destruction of gap junctions mediated by PrpL could also damage cellular cytoskeleton. The role of high pyocyanin levels in PA64481 cannot be defined in a tissue culture system *in vitro* due to the absence of serpins in this system; however, role of PrpL is emphasized because we stopped the further damage by serine protease inhibitors aprotinin and PMSF.

**Figure 3.**
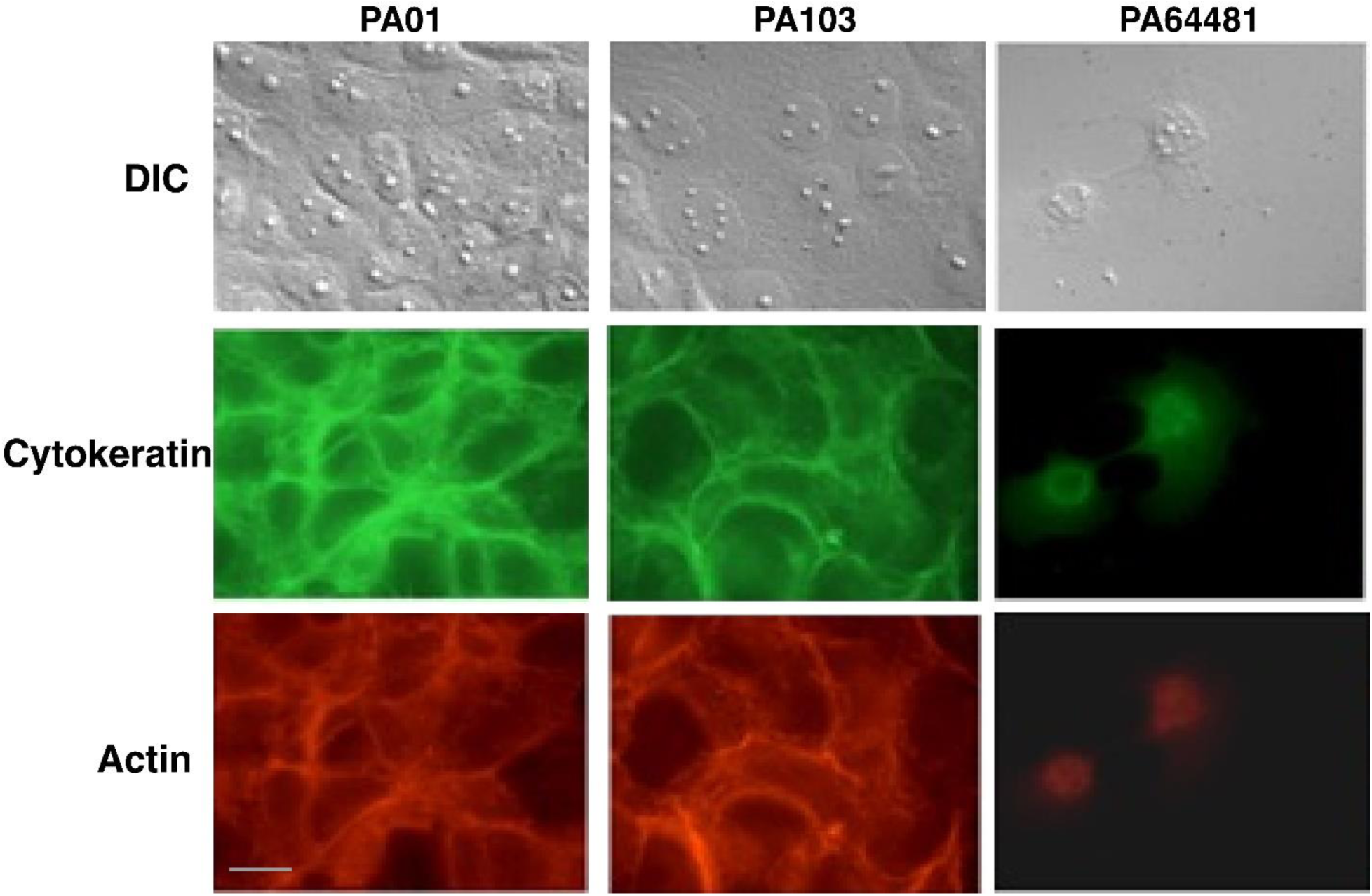
Destruction of HCE-T monolayer and cytoskeleton is associated with secreted PrpL levels in *P. aeruginosa* strains. Corneal cell monolayers on coverslips were treated with culture supernatants at 1:1 dilution in cell culture medium and were fixed at 15 minutes interval after inactivating protease by PMSF+Aprotinin. The slides were examined by Nomarski (top panels), after staining for cytokeratins followed by detection using FITC-labeled secondary antibody (middle panels), and after staining filamentous actin with TRITC-labeled phalloidin (bottom panels). Disruption of monolayers and the loss of cytoskeleton integrity can be seen within 15 minutes of treatment with supernatant from PA64481 strain, while the integrity of the cell monolayers was maintained even after 1h of treatment with both PA01 and PA103 strains supernatants. Bar indicates 10µm.

### Inactivation of type II secretion significantly reduces secretion of PrpL by *P. aeruginosa*

The halo generated by PrpL on casein plates allowed us to determine the level of secreted proteases as indicated by a “halo” around a filter disc impregnated with *P. aeruginosa* culture supernatants. We identified a Tn5Gm insertion mutant of PA64481 that did not produce a halo (Figure 4A). This mutant also did not secrete detectable level of PrpL either by silver staining or immunoblotting (Figures 4B and 4C). The observed defect was specifically attributed to the lack of secretion because high levels of PrpL was detected in bacterial lysates by Western blotting using anti-PrpL polyclonal antibodies (Figure 4D). Interestingly, the majority of PrpL protease detected by immunoblotting in bacterial lysates was found to be in mature form, i.e., migrated with an apparent molecular mass of ∼27 kD (Figure 4D). Sequencing of the DNA fragment flanking the Tn5 revealed transposon insertion into the *xcpA* gene, which encodes an essential component of the type II secretion system [31–34]. We conclude that PrpL is primarily secreted via the XcpA based type II secretion system of *P. aeruginosa*. During lysis of bacteria, cleavage by other proteases present in the extract (likely periplasmic proteases) could ultimately cleave PrpL pre-protease to produce mature enzyme. Involvement of PrpL in digestion of other secreted proteins of *P. aeruginosa* was further confirmed since protein profile of culture supernatant of *xcpA*:Tn5Gm mutant, which was comparable to that observed in PA103 and not like the parent strain PA64481, a high PrpL secreting strain (Figure 2A and 4B).

**Figure 4.**
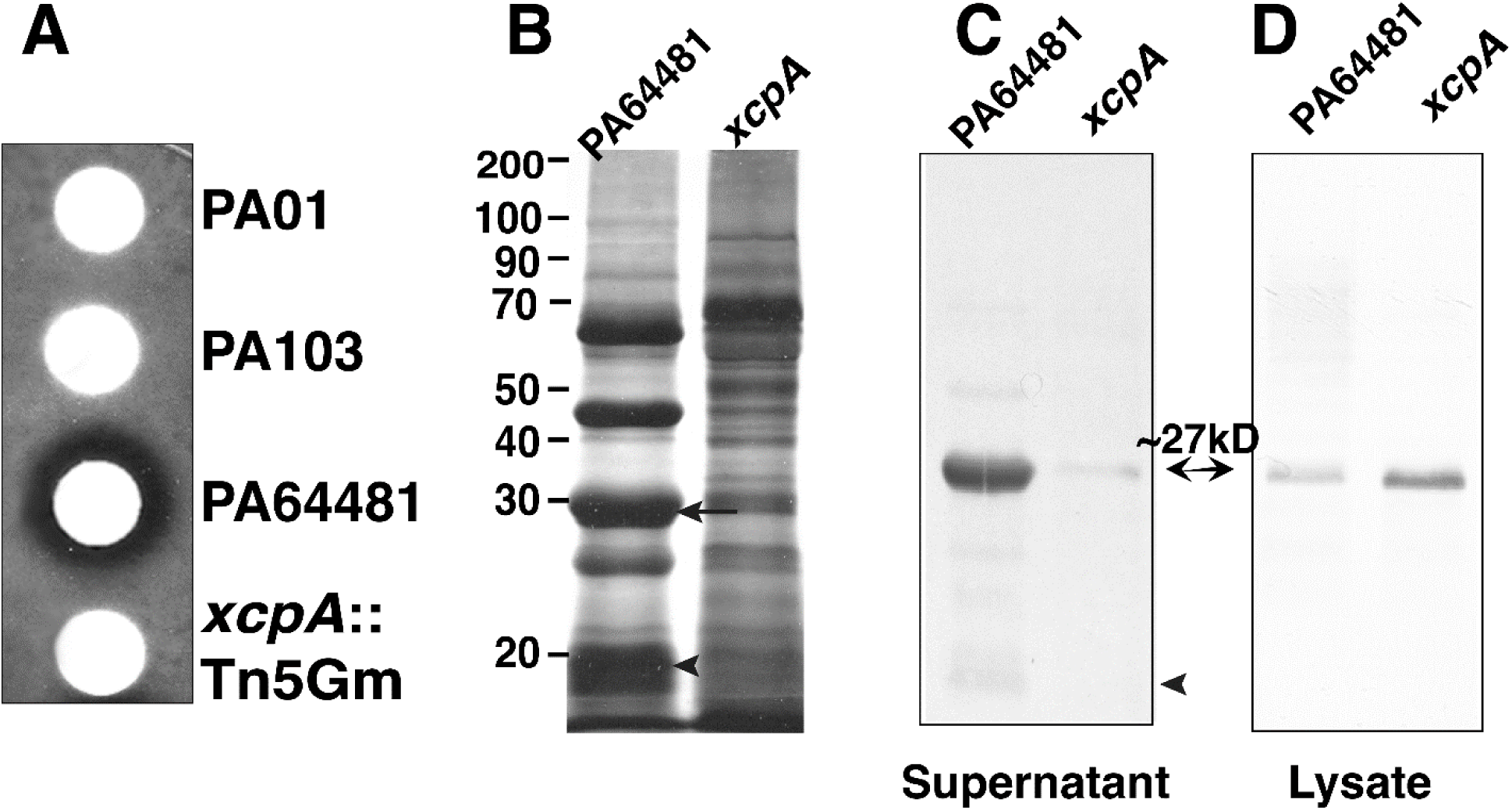
Comparative analysis of PrpL levels and activity in the *xcpA* mutant. **(A)** Filter disc caseinase assay with the culture supernatants using skim milk containing plate overlay showed no zone of clearance around the mutant in which the *xcpA* gene was disrupted by the Tn5Gm transposon and is comparable to the PA103 strain supernatant, confirming that the *xcpA* disrupted mutant fails to secrete PrpL to a significant level. **(B)** The absence of secreted protease in the culture supernatant of the *xcpA* mutant resulted in the accumulation of other secreted proteins as detected by silver staining. **(C)** Immunoblotting using anti-PrpL antiserum shows significant reduction in secreted PrpL in the *xcpA* mutant while **(D)** most of the PrpL protease remains associated with bacteria (lysate).

The results demonstrating PrpL production and secretion were further confirmed by comparing protease activity in PA64481 strain and the *xcpA* mutant culture supernatants (Figure 5A) and lysates (Figure 5B). Interestingly, supernatant from the *xcpA*:Tn5 mutant, which contains only low levels of PrpL, had only a subtle effect on the integrity of monolayers (Figure 5C) and was comparable to that observed after treatment with culture supernatants from PA01 and PA103 strains (Figure 3). Pyocyanin level in this mutant (OD_520_=1.9) was not affected.

**Figure 5.**
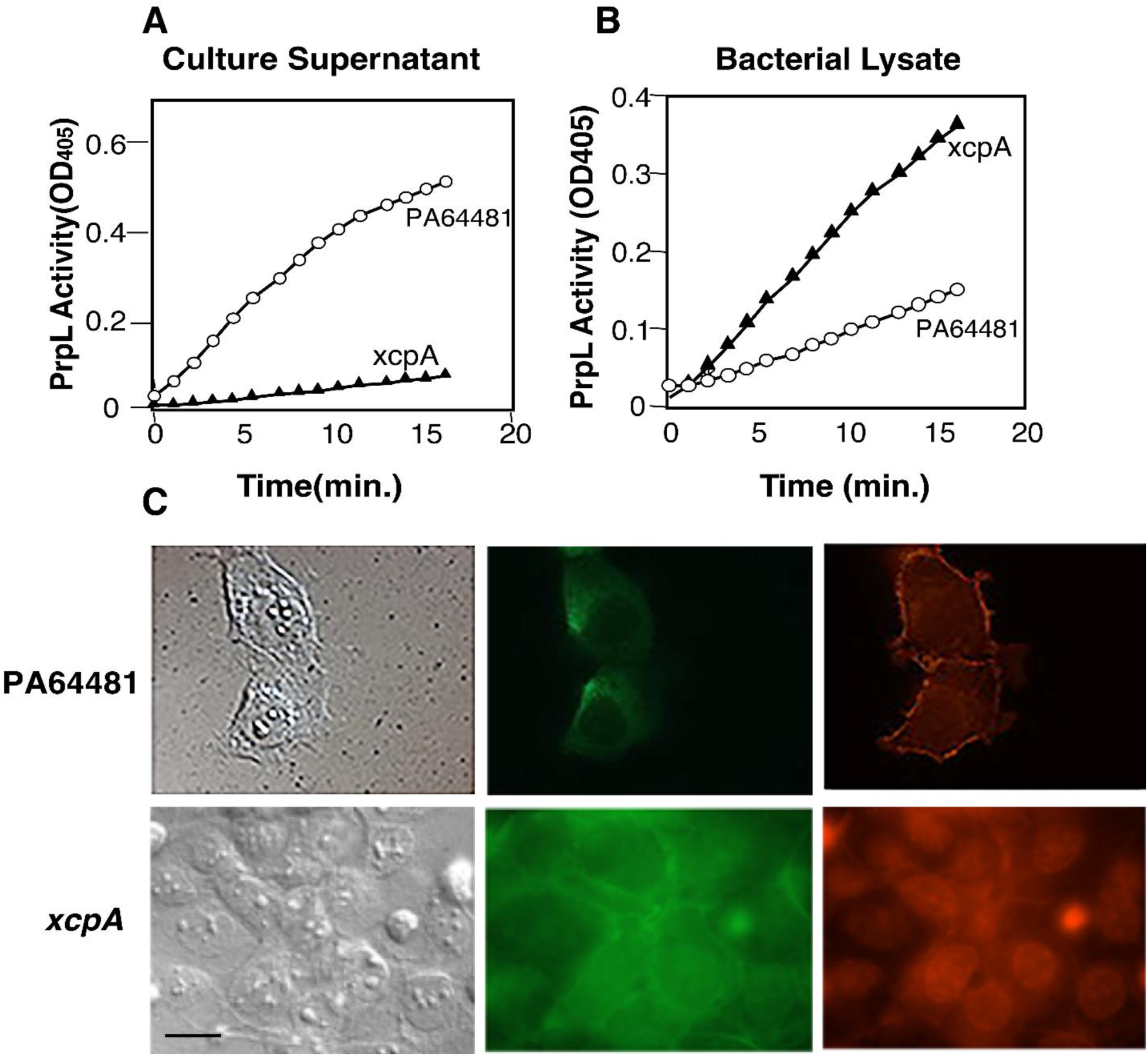
Disruption of Human Corneal Epithelial (HCE-T) cells by supernatant from high PrpL-producing strain PA64481 is eliminated in the *xcpA* mutant. (A) Minimal PrpL activity was observed in the *xcpA* mutant culture supernatant compared to that in the wild-type PA64481 strain with S2251 substrate while **(B)** significantly higher lysyl endopeptidase activity in the bacterial lysate than the parental PA64481 strain indicate failure of the mutant to secrete PrpL. **(C)** Corneal cell monolayers on coverglass were treated with culture supernatants at a 1:1 dilution in cell culture medium and were fixed at 15-minute intervals after inactivating this serine protease with PMSF+Aprotinin. The slides were examined by Nomarski (left panels), for cytokeratins using FITC-labeled secondary antibody (middle panels). Staining with TRITC-labeled phalloidin labeled filamentous actin (right panels). Disruption of monolayers and the loss of cytoskeleton can be seen within 15 minutes of treatment with supernatant from the PA64481 strain, while the integrity of the cell monolayers was maintained even after 1h treatment with the *xcpA* mutant supernatant. Bar indicates 10µm.

### Characterization of prpL knockout mutant of PA64481 strain

We could not rule out that other proteases that use Type II secretion machinery could also be involved in corneal damage. To evaluate the role of PrpL directly, we generated and characterized *prpL* knockout mutant (δPrpL) in high PrpL producing PA64481 strain (Figure 6). The absence of PrpL in culture supernatant was observed by SDS-PAGE analysis and by Western blotting. Unlike *xcpA* mutant, supernatant of the *prpL* mutant completely lacked serine protease endopeptidase activity using SS21 substrate (Figure 6C) and culture supernatant of this mutant did not disrupt HCE-T monolayers (Figure 6D).

**Figure 6.**
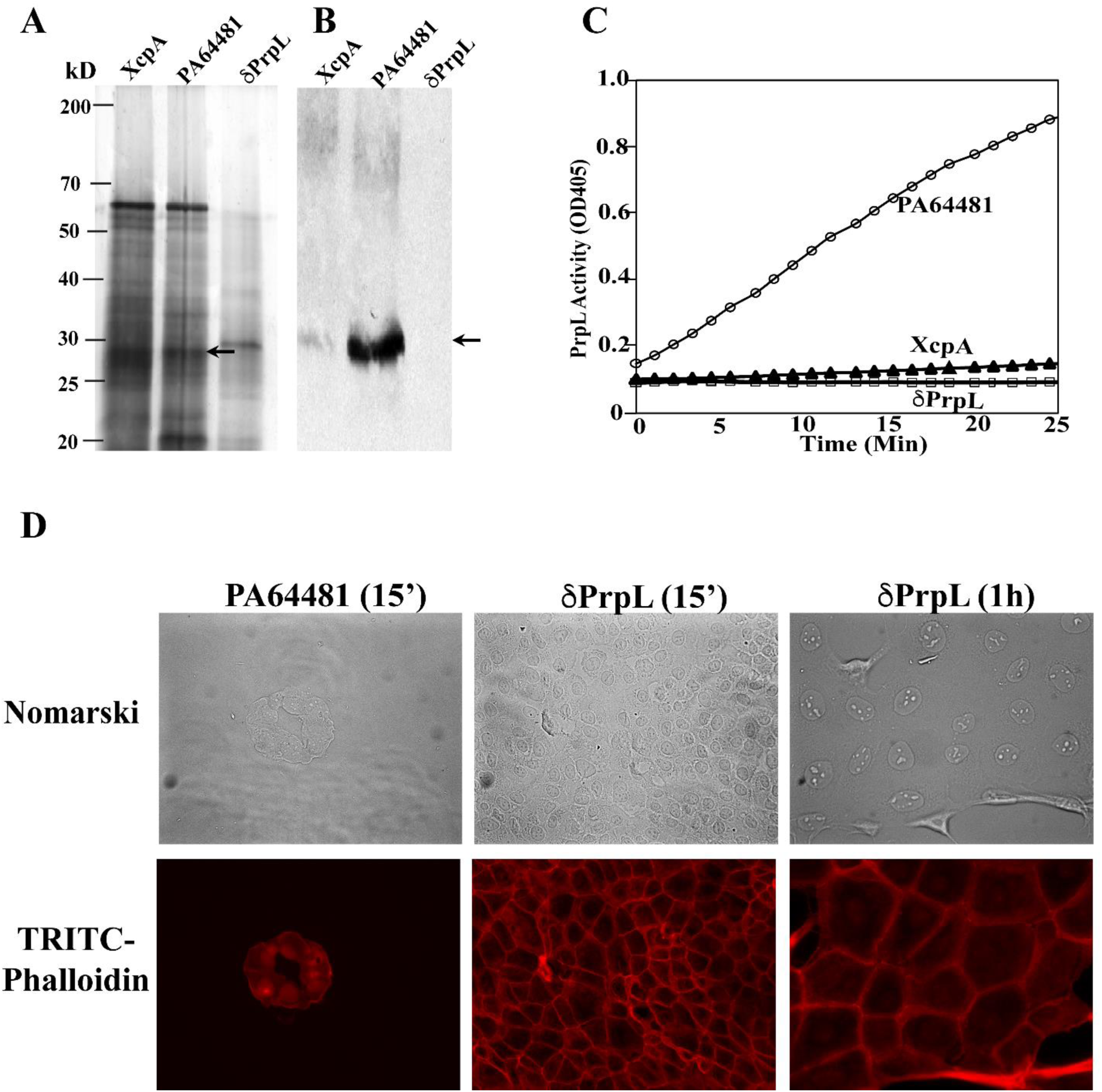
Evaluation of PrpL levels and activity in secreted milieu of the *prpL* mutant and its effect on HCE-T cell monolayer. **(A)** The absence of secreted PrpL protease in culture supernatant of the *prpL* mutant analyzed by SDS-PAGE followed by silver staining was confirmed by **(B)** Western blot analysis using anti-PrpL antiserum. **(C)** No PrpL activity was detected in the culture supernatant of the *prpL* mutant using the S2251 substrate. **(D)** After treatment of HCE-T corneal epithelial monolayer with *P. aeruginosa* culture supernatants, slides were examined by Nomarski (top panels), and after staining with TRITC-labeled phalloidin for filamentous actin (bottom panels). Unlike the wild-type PA64481 strain supernatant, secreted protein milieu of the *prpL* mutant failed to destroy the HCE-T cell monolayer even after 1h of incubation.

## Discussion

O’Callaghan and coworkers first identified PrpL (protease IV) and showed its potential role in corneal damage [13–16, 19] because a mutant lacking PrpL activity caused significantly less pathology than the parental strain in corneal scratch mouse model. Although PrpL production showed no discernible effect in a rabbit scratch model [30], severe damage in rabbit intrasomal infection model was reported as time lapsed observation after infection [14, 16]. An experimental hurdle associated with these studies is the fact that they were either performed with low PrpL producing PA103-29 strain or its derivatives.

We show here that corneal isolates of *P. aeruginosa* uniformly secrete PrpL at levels that are significantly (p=0.035) higher than that secreted by noncorneal isolates albeit there were some outliers in the latter. In our assays, PA103 supernatant demonstrated no lysyl endopeptidase activity, even when the assays were extended to 1h, suggesting exquisite sensitivity of the substrate to PrpL protease. A previous study with a strain with high PrpL activity did not address its possible role in tissue damage or pathogenesis [35].

We show that PA64481 secretes large amounts of PrpL with potent catalytic activity and significantly higher level of pyocyanin. An insertion in the *xcpA* gene of PA64481 abolished secretion of PrpL but not its expression. Given that XcpA is a peptidase involved in processing of proteins that are suggested to form the type II secretory machinery in the outer membrane of *P. aeruginosa* [31–34], our results suggest that PrpL is secreted primarily via this type II secretion machinery similar to lipase and elastases, which are also secreted by the same system [31]; however, PrpL and not elastase is usually expressed during *P. aeruginosa* infection [36] and thus, may not play a major role in corneal damage [12, 37–39] or damage to collagen by cleavage and activation of the host pro-matrix metalloproteinases during infection [40].

Corneal damage is a hallmark of serious infections by *P. aeruginosa*, and the activity of PrpL likely contributes to epithelium destruction significantly. Consistent with this notion, we show here that the ability to damage monolayers of corneal epithelial cells is associated with the secretion of PrpL at high levels and the absence of PrpL fails to disrupt HCE-T corneal epithelial cell monolayer observed by the culture supernatant of the wild-type PA64481 strain (Figures 5 and 6). Our kinetic analysis using S2251 substrate of the *prpL* mutant supernatant established that this is the only secreted serine protease in *P. aeruginosa*.

A role for PrpL may not be restricted to corneal infections because we show here that several non-corneal *P. aeruginosa* isolates expressed detectable levels of lysyl endopeptidase activity. In a previously conducted competition experiments with wild type PA01 and a PA01-derived disruption mutant implicated PrpL in the efficient colonization of rat lungs [21]. It is easy to imagine that higher PrpL protease activity, as well as other secreted *P. aeruginosa* proteases, contributes to the pulmonary injury suffered by patients with acute pneumonia or cystic fibrosis-associated lung disease. Identification of *P. aeruginosa* clinical isolates that produce high levels of PrpL and pyocyanin will facilitate future studies involving evaluation of the role of these molecules in colonization and damage of various tissues during infection. Furthermore, the characterization of various regulatory mechanisms that contribute to the production of both PrpL and pyocyanin together, or separately, could help identify overall pathogenesis mechanisms of *P. aeruginosa* and damage in corneal and wound infections in mammalian hosts as incurred by these virulence factors individually, or together.

## ACKNOWLEDGEMENTS

This study was supported by the faculty accounts of JDG and NP.

