## Supplementary Information for "Retrospective analysis of *Pseudomonas aeruginosa* clinical isolates for secreted PrpL protease: a key virulence factor associated with corneal tissue damage"

**SUPPLEMEMTARY INFORMATION.**

**Methods.**

Primers based on the known sequence of *prpL* of *P. aeruginosa* strain PA01 [38], 5’ ATGCATAAGAGAACGTACCTGAATG 3’ and 5’TCAGGGCGCGAAGTAGCGGGA 3’ and Hercules polymerase (Stratagene Co. Inc.) were used to amplify, clone and sequence the *prpL* genes from *P. aeruginosa* PA64481, PA01 and PA103. The PCR products were cloned in Topo-XL vector (Invitrogen Inc.) and sequenced. Among 68 strains evaluated in this study, six strains genomes sequencing have been completed with Accession numbers for three major strains: PA01 (QZFY00000000), PA103 (NZ_JAMDFM000000000), PA14 (AY273869).

**Supplementary Figure 1A. PrpL protein sequence alignment of three *P. aeruginosa* strains depicting signal peptide (blue), N-terminal sequencing of the mature protein (green), and active site triad residues: His, Asp, Ser (magenta) are marked. Percent identity matrix is shown below.**

```
PA01         MHKRTYLNACLVLALAAGASQALAAPGASEMAGDVAVLQASPASTGHARFANPNAAISAA	60
PA103        MHKRTYLNACLVLALAAGASQASAAPGASEMAGDVAVLQASPASTGHARFANPNAATSAA	60
PA64481      MHKRTYLNACLVLALAAGASQALAAPGASEMAGDVAVLQASPASTGHARFANPNAAISAA	60
             ********************** ********************************* ***

PA01         GIHFAAPPARRVARAAPLAPKPGTPLQVGVGLKTATPEIDLTTLEWIDTPDGRHTARFPI	120
PA103        GIHFAAPPARRVARAAPLAPKPGTPLQVGVGLKTATPEIDLATLEWIDTPDGRHTARFPI	120
PA64481      GIHFAAPPARRVARAAPLAPKPGTPLQVGVGLKTATPEIDLTTLEWIDTPDGRHTARFPI	120
             *****************************************:******************

PA01         SAAGAASLRAAIRLETHSGSLPDDVLLHFAGAGKEIFEASGKDLSVNRPYWSPVIEGDTL	180
PA103        SAAGAASLRAAIRLETRSGSLPDDVLLHFAGAGKEIFEASGKDLSLNRPYWSPVIEGDTL	180
PA64481      SAAGAASLRAAIRLETRSGSLPDDVLLHFAGAGKEIFEASGKDLSVNRPYWSPVIEGDTL	180
             ****************:****************************:**************

PA01         TVELVLPANLQPGDLRLSVPQVSYFADSLYKAGYRDGFGASGSCEVDAVCATQSGTRAYD	240
PA103        TVELVLPANLQPGDLRLSVPQVSYFADSLYKAGYRDGFGASGSCEVDAVCATQSGTRAYD	240
PA64481      TVELVLPANLQPGDLRLSVPQVSYFADSLYKAGYRDGFGASGSCEVDAVCATQSGTRAYD	240
             ************************************************************

PA01         NATAAVAKMVFTSSADGGSYICTGTLLNNGNSPKRQLFWSAAHCIEDQATAATLQTIWFY	300
PA103        NATAAVAKMVFTSSADGGSYICTGTLLNNGNPPKRQLFWSAAHCIEDQATAATLQTIWFY	300
PA64481      NATAAVAKMVFTSSADGGSYICTGTLLNNGNSPKRQLFWSAAHCIEDQATAATLQTIWFY	300
             ******************************* ****************************

PA01         NTTQCYGDASTINQSVTVLTGGANILHRDAKRDTLLLELKRTPPAGVFYQGWSATPIANG	360
PA103        NTTQCYGDASTINQSVTVLTGGANILHRDAKRDTLLLELKRTPPAGVFYQGWSATPIANG	360
PA64481      NTTQCYGDASTINQSVTVLTGGANILHRDAKRDTLLLELKRTPPAGVFYQGWSATPIANG	360
             ************************************************************

PA01         SLGHDIHHPRGDAKKYSQGNVSAVGVTYDGHTALTRVDWPSAVVEGGSSGSGLLTVAGDG	420
PA103        SLGHDIHHPRGDAKKYSQGNVSAVGVTYDGHTALTRVDWPSAVVEGGSSGSGLLTVAGDG	420
PA64481      SLGHDIHHPRGDAKKYSQGNVSAVGVTYDGHTALTRVDWPSAVVEGGSSGSGLLTVAGDG	420
             ************************************************************

PA01         SYQLRGGLYGGPSYCGAPTSQRNDYFSDFSGVYSQISRYFAP------------------	462
PA103        SYQLRGGLYGGPSYCGAPTSQRNDYFSDFSGVYSQISRYFAP------------------	462
PA64481      SYQLRGGLYGGPSYCGAPTSQRNDYFSDFSGVYSQISRYFAP------------------	462
             ******************************************
```

**Percent Identity Matrix**

```
PA01		100.00%	98.70%	99.78%
PA103		98.70%	100.00%	98.92%
PA64481	98.92%	99.78%	100.00%
```

**Supplementary Figure 1B. PrpL of *P. aeruginosa* is slightly more closely related by amino acid sequence to *Lysobacter enzymogenes* arginal endopeptidase, LeR than lysyl endopeptidase of the same species.**

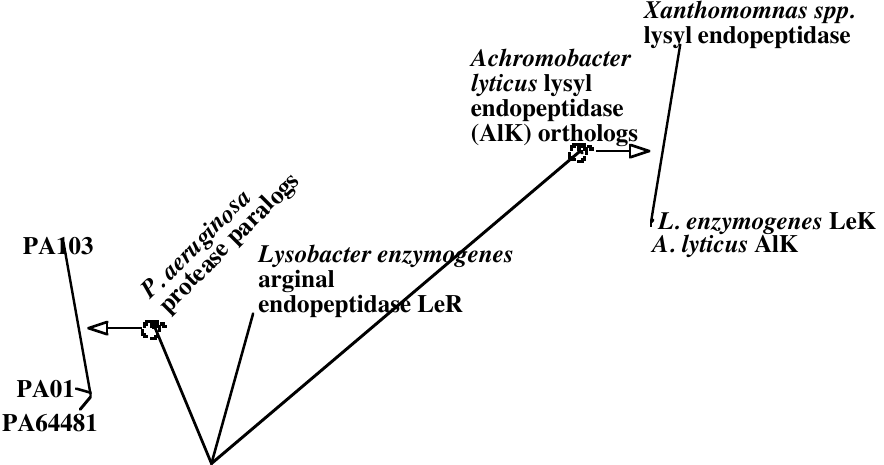
